# spotPCR: A Rapid and Efficient Approach for Indexing Individual Template Molecules using Unique Molecular Identifiers

**DOI:** 10.1101/2024.12.11.627791

**Authors:** Jason D Limberis, Roland J Nagel, Soumitesh Chakravorty, Dena Block, Scot Dewell, Alina Nalyvayko, John Z Metcalfe

## Abstract

Low-frequency mutations provide valuable insights in various fields, including drug resistance identification, cancer and infectious disease research. One promising strategy to enhance the sensitivity and specificity of mutation detection is the incorporation of unique molecular identifiers (UMIs) during polymerase chain reaction (PCR) amplification and before deep sequencing. However, conventional methods for UMI incorporation oRen necessitate multiple labor-intensive steps. spotPCR (Specific Primer Limited Unique Molecular Identifier Tagging PCR) overcomes these challenges, streamlining the UMI tagging process.

## Introduction

Low-frequency mutations provide valuable insights in various fields. In cancer research, they contribute to our understanding of tumor heterogeneity and clonal evolution; in infectious diseases, they allow us to identify drug resistance and track transmission patterns; and in personalized medicine, they assist us in predicting treatment response and optimizing therapeutic strategies^1,2^. One promising strategy to enhance the sensitivity and specificity of mutation detection is the incorporation of unique molecular identifiers (UMIs) before polymerase chain reaction (PCR) amplification and deep sequencing. UMIs are molecular barcodes that distinguish true mutations from PCR or sequencing errors by identifying consensus between reads that share the same barcode. They also allow for the identification of PCR duplicates and therefore the more accurate counting of molecules, and the use of non-high fidelity but robust polymerases. However, conventional methods for UMI incorporation often necessitate several labor-intensive steps, such as multiple PCRs separated by clean-ups, ligations, and exonuclease digestions^3^. This is because molecules must first be tagged with UMIs, often by ligating UMI containing Illumina adapters, and then diluted to a level that can then be amplified and sequenced to allow for several sequencing reads per initially tagged molecule (i.e., the UMI family). These steps increase the complexity, time, and cost associated with library preparation, thereby limiting the widespread adoption of UMIs for mutation detection.

To overcome these challenges, we present an innovative approach termed spotPCR (Specific Primer Limited Unique Molecular Identifier Tagging PCR; Fig. 1), which offers a rapid, single-tube nested PCR-based method to generate sequenceable-ready libraries containing UMIs. SPOT-PCR streamlines the UMI tagging process, removing the need for multiple enzymatic treatments, and simplifying the workflow. SPOT-PCR adds UMIs to individual molecules using a limiting amount of target-specific primers containing a UMI and a universal tail sequence. The universal tail sequences are then used to amplify the sequencing library.

**Fig 1.**
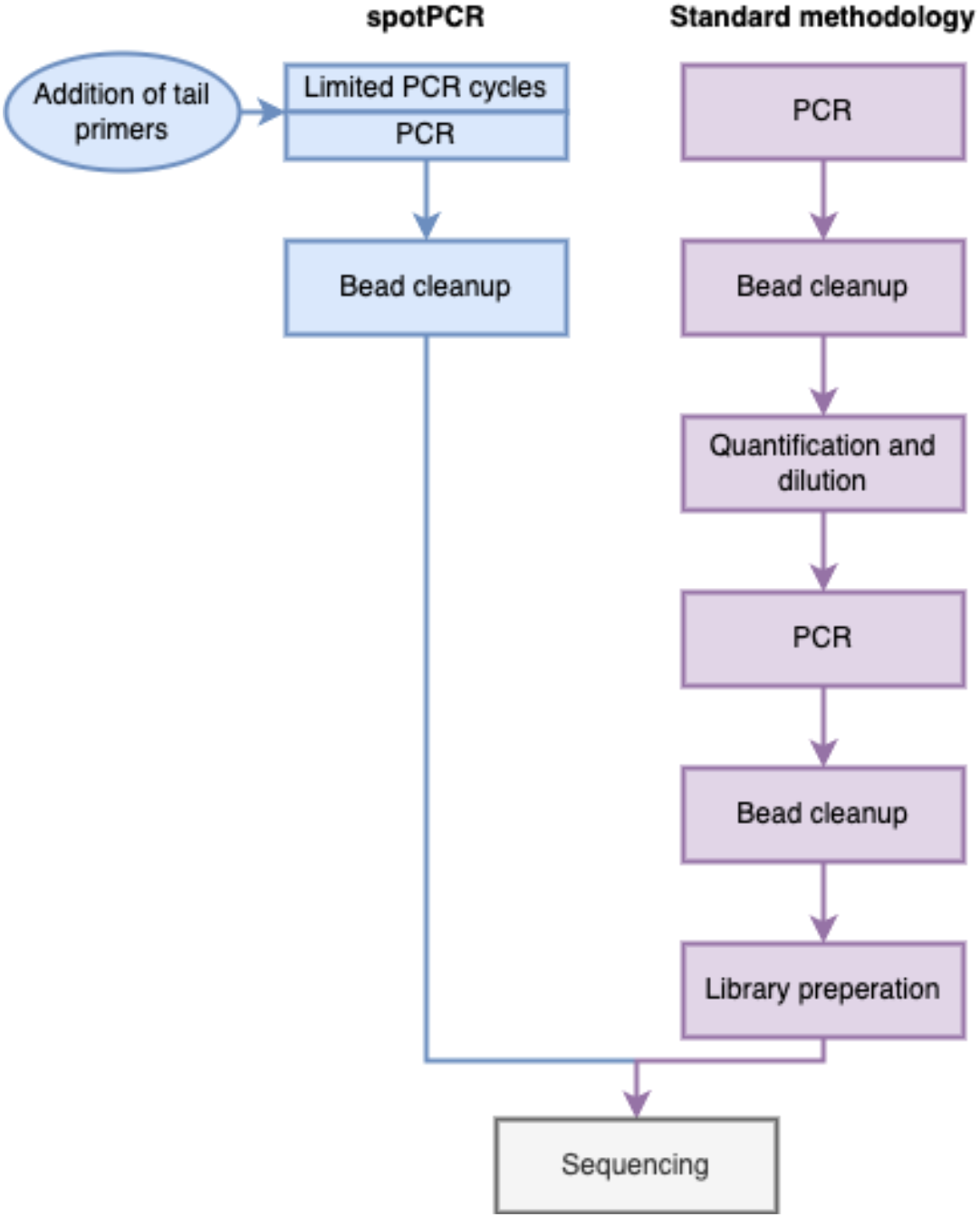
spotPCR workflow is much simpler than a standard, UMI tagging approach.

## Materials and Methods

The protocol described is published on protocols.io, DOI 10.17504/protocols.io.261ged66wv47/v1.

## Expected Results

After obtaining a band of the expected size with a TapeStation (Agilent, USA) or an agarose gel, the prepared library can be quantified using the appropriate kit for the chosen sequencing method (e.g., the NEBNext Library Quant Kit for Illumina [NEB, USA]) to provide accurate concentrations of the sequenceable amplicons (as not all the amplicons generated will have both adapters and be sequenceable, thus, global quantification methods like Qubit [Applied Biosystems, USA] will overestimate the sequenceable quantity). The libraries are then sequenced according to the manufacturer’s instructions. The provided pipeline processes Illumina sequencing data, identifies the UMIs in the reads, maps the reads to a reference, and calls mutations.

In this example, we did a duplex spotPCR using different *limiting* primer concentrations on a single UMI-containing primer (RYNNNRYNNNTT) targeting two drug resistance-conferring regions, *gyrA* and *pncA*, of the *Mycobacterium tuberculosis* genome, followed by 150bp Illumina pair-end sequencing. **Table 1** presents various sequencing statistics and UMI identification output by the pipeline, demonstrating that at an initial limiting primer amount of 0.0625uM for both *gyrA* and *pncA* provides usable UMI-tagged reads for most reads while other concentrations provide less usable UMI-tagged reads. The efficiency of the *pncA* primer set is lower than that of the *gyrA* set, hence it reaches a high UMI tagging proportion even at high limiting primer concentrations.

**Table 1.**
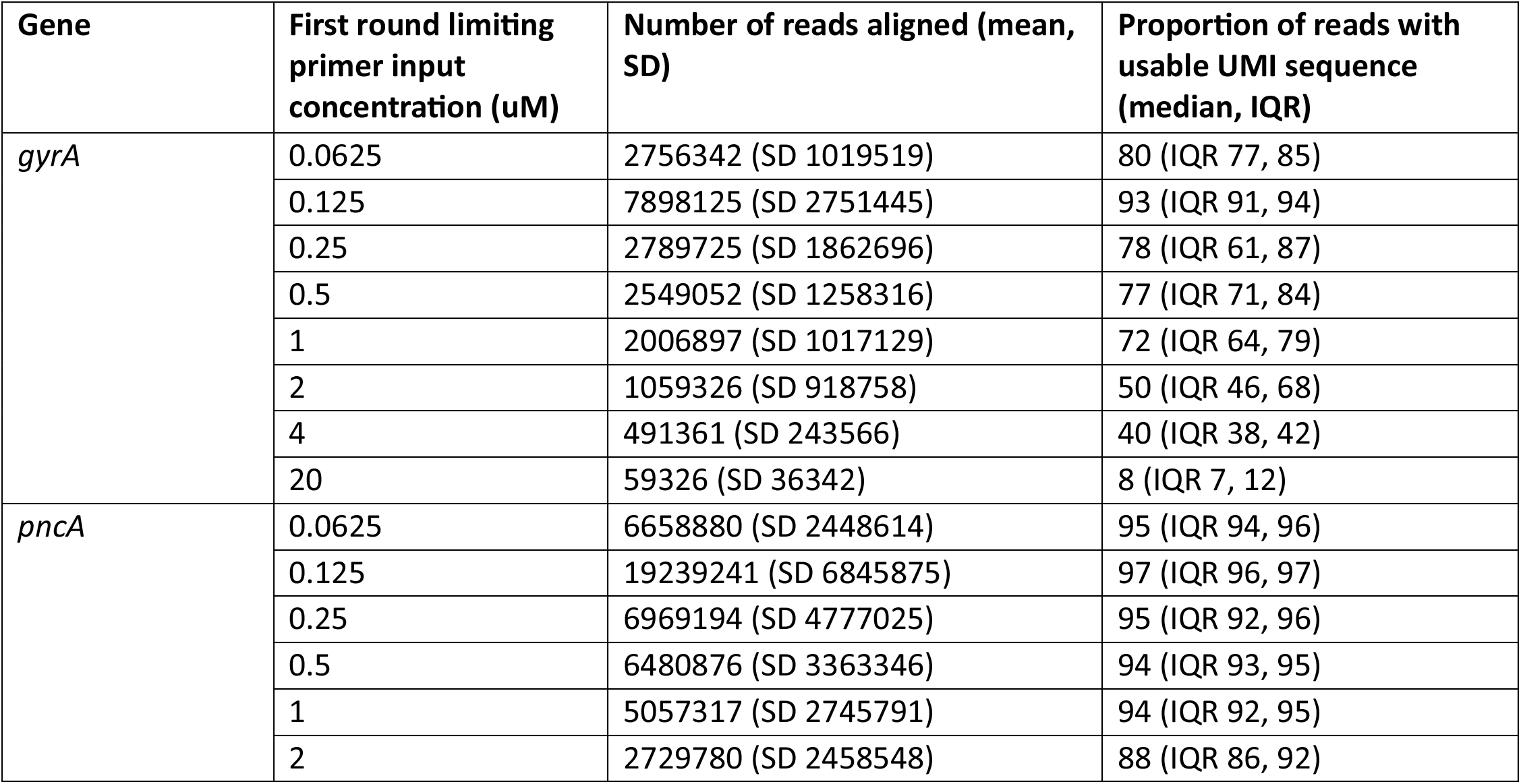
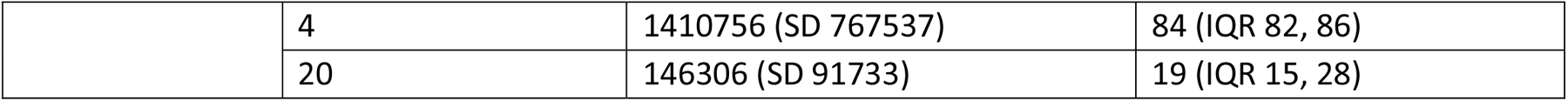
Sequencing and UMI statistics filtering for UMIs occurring fifty or more times for a duplex spotPCR of two *M. tuberculosis* genes at differing limiting first-round primer concentrations with six replicates per group shows that a primer concentration of 0.125 and 0.0625 allows for successful spotPCR UMI tagging for both targets.

## Supporting information

Protocol S1

## Supporting Information

Sequence data is available on BioProject under the accession number PRJNA1176610.

### CompeAng interests

RJN, DB, SD, and SC are Cepheid employees. Other authors have no competing interests.

