## Supplementary material for "spotPCR: A Rapid and Efficient Approach for Indexing Individual Template Molecules using Unique Molecular Identifiers": Protocol S1

Jason D Limberis[1]

[1]University of California, San Francisco

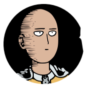

Jason D Limberis

University of California, San Francisco

**Protocol Info:** Jason D Limberis . spotPCR: A Rapid and Efficient Approach for Indexing Individual Template Molecules using Unique Molecular Identifiers. **protocols.io** **https://protocols.io/view/spotpcr-a-rapid-and-efficient-approach-for-indexin-cvuaw6se**

**Created:** June 16, 2023

**Last Modified:** September 16, 2024

**Protocol Integer ID:** 83554

### Materials

**Required**

Q5 Hot Start High-Fidelity DNA Polymerase - 500 units **New England Biolabs** **Catalog #**M0493L

Agencourt AmPure XP beads **Contributed by users** **Catalog #**A63880

dNTPs **Contributed by users**

A thermocycler and a qPCR machine

A magnetic rack

Optional

NEBNext Library Quant Kit for Illumina - 500 rxns **New England Biolabs** **Catalog #**E7630L

| A | B | C |
|---|---|---|
| Primer Set | Direction | Sequence |
| pncA | F | TCGTCGGCAGCGTCAGATGTGTATAAGAGACAGCGCGGCGTCATGGACCCTAT |
| pncA | R | GTCTCGTGGGCTCGGAGATGTGTATAAGAGACAGNNNNNNNNNNTTTCGAAGCCGCTGTACGCTCC |
| gyrA | F | TCGTCGGCAGCGTCAGATGTGTATAAGAGACAGTCACCCGCAACGCCAAGGAT |
| gyrA | R | GTCTCGTGGGCTCGGAGATGTGTATAAGAGACAGNNNNNNNNNNTTATTGCCTGGCGAGCCGAAGT |
| Illumina adapter primer | F | CAAGCAGAAGACGGCATACGAGAT[i7]GTCTCGTGGGCTCGGAGATGTGTATAAGAGACAG |
| Illumina adapter primer | R | AATGATACGGCGACCACCGAGATCTACAC[i5]TCGTCGGCAGCGTCAGATGTGTATAAGAGACAG |

You can use any compatable Illumina adapters such as the IDT for Illumina DNA/RNA UD Indexes.

#### Stage 1 PCR

1

| A | B |
|---|---|
| COMPONENT | Volume (µl) |
| 5X Q5 Reaction Buffer | 5 |
| 5X Q5 High GC Buffer | 5 |
| 10 mM dNTPs | 0.5 |
| Q5 High-Fidelity DNA Polymerase | 0.25 |
| 1µM Forward primer 1 | 1 |
| 1µM Forward primer 2 | 1 |
| 0.0625µM Reverse primer 1* | 1 |
| 0.625µM Reverse primer 2* | 1 |
| Template DNA | 2 |
| Nuclease-Free Water | 6.25 |

*We recommend doing a dilution series starting at 500fM to asses new primersets.

| A | B | C | D |
|---|---|---|---|
| Step | Temp (C) | Time (s) | Cycles |
| Denaturation | 98 | 120 | 1 |
| Denaturation | 98 | 10 | 3 |
| Annealing | 62 | 45 | |
| Extension | 72 | 120 | |
| Extension | 4 | Forever | 1 |

Cycle parameters

#### Stage 2 PCR

2 Add the below into the reaction while at 4 °C and proceed to the second PCR cycling

| A | B |
|---|---|
| COMPONENT | Volume (µl) |
| Illumina adapter primer F (10uM) | 1 |
| Illumina adapter primer R (10uM) | 1 |

| A | B | C | D |
|---|---|---|---|
| Step | Temp (C) | Time (s) | Cycles |
| Denaturation | 98 | 120 | 1 |
| Denaturation | 98 | 5 | 34 |
| Annealing | 65 | 15 | |
| Extension | 72 | 20 | |
| Extension | 4 | Forever | 1 |

Cycle parameters

#### Bead cleanup

21m

3 Add 25 µL (1X) of resuspended AMPure XP Beads to the sample

Mix by pipetting 10x

4 Incubate 00:02:00 at Room temperature

2m

5 Place on the magnet, allow the beads to aggregate, and remove and discard the supernatant

6 Add 200 µL [M] 70 % (v/v) ethanol and incubate (still on the magnet) for 00:00:30

30s

6.1 Remove the supernatant

6.2 Repeat go to step #6 for a total of 2 washes

7 Air dry for 00:00:30 , don't allow the beads to become cracked

30s

8 Immediately after the bead pellet becomes opaque, remove the tube from magnetic rack and resuspend in 20 µL of **Low EDTA Tris Buffer**. Ensure all beads are in solution.

9 Incubate at room temperature for 00:05:00

10 Place on magnetic rack, wait for the solution to become clear ~ 00:02:00 , and transfer the eluted DNA to a new well-labeled tube

#### Optional: NEB Illumina Quantification

11 Thaw the NEBNext Library Quant Master Mix and NEBNext Library Quant Primer Mix. Ensure mixing of NEBNext Library Quant Primer Mix by vortexing. Place reagents on ice.

12 Thaw the NEBNext Library Quant DNA Standards, tubes 1–6.
Mix by pulse vortexing on a low setting. Briefly spin to collect material from the sides of the tubes. Place on ice.

13 Thaw the NEBNext Library Quant Dilution Buffer (10X). Mix well by vortexing.
Centrifuge briefly to collect material from the sides of the tube.
Place on ice.

14 Add 100 µL NEBNext Library Quant Primer Mix to the tube of NEBNext Library Quant Master Mix ( 1.5 mL ). Mix by vortexing. Write the date on the master mix tube to indicate that primer mix has been added.

15 Dilute the NEBNext Library Quant Dilution Buffer (10X) 1:10 with nuclease-free water. Mix by vortexing.
Prepare sufficient buffer for quantitating the desired number of libraries, allowing 1.2 mL for each library.

16 Prepare a 1:1,000 dilution of each library sample in NEBNext Library Quant Dilution Buffer (1X)

17 Aliquot 16 µL NEBNext Library Quant Master Mix (with primers) to each well

18 Add-in 4 µL of sample or standard per well

19

| A | B | C | D |
|---|---|---|---|
| Step | Temp (C) | Time (s) | Cycles |
| Denaturation | 95 | 60 | 1 |
| Denaturation | 95 | 15 | 35 |
| Annealing | 63 | 45 | |

Cycle parameters

A denaturation/melt curve can be included if desired, but is optional.

| A | B |
|---|---|
| Sample | Conc. (pM) |
| DNA Standard 1 | 100 |
| DNA Standard 2 | 10 |
| DNA Standard 3 | 1 |
| DNA Standard 4 | 0.1 |
| DNA Standard 5 | 0.01 |
| DNA Standard 6 | 0.001 |

20 Adjusted Conc. = Calculated Conc. × 399 / library size (bp)

#### Analysis

21 **Link to script (click here)**

```
bash spotPCR_process.sh \
  -R1 "Read1.fastq" \
  -R2 "Read2.fastq" \
  -Ref_name "Reference.fasta" \
  -sample_name "SampleName" \
  -threads "Threads" \
  -umi "UMI_Barcode_Pattern"
```
